# Intra-session test-retest reliability of functional connectivity in infants

**DOI:** 10.1101/2020.06.25.169524

**Authors:** Yun Wang, Walter Hinds, Cristiane S Duarte, Seonjoo Lee, Catherine Monk, Melanie Wall, Glorisa Canino, Ana Carolina C. Milani, Andrea Jackowski, Marina Griorgi Mamin, Bernd U. Foerster, Jay Gingrich, Myrna M Weissman, Bradley S. Peterson, David Semanek, Edna Acosta Perez, Eduardo Labat, Ioannisely Berrios Torres, Ivaldo Da Silva, Camila Parente, Nitamar Abdala, Jonathan Posner

**Affiliations:** Department of Psychiatry, Columbia University Medical Center, New York, NY, USA; Mental Health Data Science, New York State Psychiatric Institute, New York, NY, USA; Department of Obstetrics and Gynecology, New York State Psychiatric Institute, New York, NY, USA; School of Medicine, University of Puerto Rico, San Juan, PR, USA; Department of Psychiatry, Federal University of Sao Paulo, Sao Paulo, Brazil; Interdisciplinary Lab for Clinical Neurosciences, Federal University of Sao Paulo, Sao Paulo, Brazil; Department of Pediatric, Federal University of Sao Paulo, Sao Paulo, Brazil; Institute for the Developing Mind, The Saban Research Institute, Children’s Hospital Los Angeles, CA, USA; Department of Gynecology, Federal University of Sao Paulo, Sao Paulo, Brazil; Department of Diagnostic Radiology, Federal University of Sao Paulo, Brazil

**Keywords:** Infant, fMRI, Test-retest reliability, ICC, Edge-level, Subject-level, Network, Jaccard Index, ICA

## Abstract

Resting functional MRI studies of the infant brain are increasingly becoming an important tool in developmental neuroscience. Whereas the test-retest reliability of functional connectivity (FC) measures derived from resting fMRI data have been characterized in the adult and child brain, similar assessments have not been conducted in infants. In this study, we examined the intra-session test-retest reliability of FC measures from 119 infant brain MRI scans from four neurodevelopmental studies. We investigated edge-level and subject-level reliability within one MRI session (between and within runs) measured by the Intraclass correlation coefficient (ICC). First, using an atlas-based approach, we examined whole-brain connectivity as well as connectivity within two common resting fMRI networks – the default mode network (DMN) and the sensorimotor network (SMN). Second, we examined the influence of run duration, study site, and scanning manufacturer (e.g., Philips and General Electric) on ICCs. Lastly, we tested spatial similarity using the Jaccard Index from networks derived from independent component analysis (ICA). Consistent with resting fMRI studies from adults, our findings indicated poor edge-level reliability (ICC = 0.14 - 0.18), but moderate-to-good subject-level intra-session reliability for whole-brain, DMN, and SMN connectivity (ICC = 0.40 - 0.78). We also found significant effects of run duration, site, and scanning manufacturer on reliability estimates. Some ICA-derived networks showed strong spatial reproducibility (e.g., DMN, SMN, and Visual Network), and were labelled based on their spatial similarity to analogous networks measured in adults. These networks were reproducibly found across different study studies. However, other ICA-networks (e.g. Executive Control Network) did not show strong spatial reproducibility, suggesting that the reliability and/or maturational course of functional connectivity may vary by network. In sum, our findings suggest that developmental scientist may be on safe ground examining the functional organization of some major neural networks (e.g. DMN and SMN), but judicious interpretation of functional connectivity is essential to its ongoing success.

**Highlights:**

- Infant functional connectivity (FC) shows poor edge-level reliability (ICCs)
- However, subject-level infant FC estimates show good-to-excellent ICCs
- Spatial reproducibility is better for some resting networks (DMN, SMN) than others (ECN)
- Reliability estimates differ across study site and MRI scanner
- Conclusion - Infant FC can be a reliable measurement, but judicious use is needed

## 1. INTRODUCTION

Providing a window into early brain development with limited influence of the post-natal environment, resting functional MRI (fMRI) studies of the infant brain have become an important tool for advancing developmental neuroscience (Duarte et al., 2020; Posner et al., 2016). Subtle changes in brain function early in development may, for example, confer lifelong implications on cognition, behavior, and psychopathology. Resting fMRI studies detail the organization of neural circuits based on functional connectivity, or the coherence of neural activity across disparate brain regions. In infant research, this approach may help identify early brain-based biomarkers, providing, for example, indicators of aberrant neurodevelopment, prenatal exposure effect, or neurobiological correlates of early cognitive functions, and may thereby facilitate early identification of at-risk individuals (Lugo-Candelas et al., 2018; Posner et al., 2016; Rudolph et al., 2018). Despite this promise and the growing use of resting fMRI studies in infant research, the reliability of functional connectivity measures in the infant brain remains largely unexamined. Establishing test-retest reliability is critical to solidifying the methodological rigor and basis for including resting fMRI techniques within the armamentarium of neurodevelopmental researchers.

Prior studies have examined the intra-session (so-called short-term) and between-session test-retest reliability of functional connectivity measures in the adult and child brain (Marchitelli et al., 2016; Noble et al., 2019; Somandepalli et al., 2015). Most studies have used the Intraclass correlation coefficient (ICC) as the primary measure of test-retest reliability for functional connections due to the flexibility of the ICC to explicitly model known sources of measurement variance (Shrout and Fleiss, 1979; Somandepalli et al., 2015). Across a wide range of approaches, these studies have typically found poor reliability for edge-level ICCs (Noble et al., 2019), yet moderate-to-good reliability for subject-level ICCs (Mejia et al., 2018; Mejia et al., 2015; Somandepalli et al., 2015). In the context of reliability for functional connectivity, an “edge-level” ICC refers to the test-retest reliability of a single functional connection (an edge). For every pairwise set of edges between regions-of-interest (ROIs) across the brain, an ICC can be calculated resulting in thousands of ICCs per brain scan. In contrast, a “subject-level” ICC is calculated across a connectivity matrix – that is, multiple connections within one brain scan, or within one brain network, are considered collectively.

Because of the temporally correlated nature of time series fMRI data, the time interval between scanning sessions (i.e., short-term versus long-term) is an important consideration when assessing reliability. Several studies, for example, have found repeated measures taken over shorter intervals (e.g. intra-session or within days) are more reliable than those taken over longer intervals (e.g. weeks to years). Herein, we examine intra-session test-retest reliability for two primary reasons. First, intra-session reliability has not yet been established in infants. Second, long-term reliability in infants may be confounded by neurodevelopmental changes that occur rapidly in infancy.

Resting fMRI reliability estimates may be influenced by MRI acquisition and processing procedures. For example, studies have reported differences in connectivity depending on the MRI scanning site (Biswal et al., 2010; Jovicich et al., 2016). These effects are detected even after harmonization techniques are conducted (Noble et al., 2017). Also, reliability improves with the acquisition of more frames of fMRI data per subject either by increasing the durations of the scanning sessions or by increasing the number of runs (Birn et al., 2013). Lastly, several studies have found that pre-processing procedures designed to reduce the contribution of noise, such as motion, global signal, white matter, and cerebrospinal fluid, to fMRI signal can have unexpected effects on reliability estimation (Guo et al., 2012; Parkes et al., 2018; Shirer et al., 2015). For example, motion correction, a technique used to reduce the influence of head motion on fMRI signal, can, somewhat counterintuitively, *reduce* test-retest reliability when the motion itself is consistent over time (Parkes et al., 2018). Of note, ROIs from different atlases can vary substantially (Salehi et al., 2020), resulting in different ICCs.

Despite the numerous test-retest reliability studies that have been conducted in adults and children, prior studies have not reported reliability estimates in infants, nor have they examined test-retest reliability across infant studies conducted at different scanning sites. The moderate-to-good reliability for subject-level ICCs reported in studies of children and adults (e.g.(Marchitelli et al., 2016; Noble et al., 2017; Somandepalli et al., 2015)) is not sufficient to assume similar reliability will also be present in infants. First, significant neurovascular changes occur throughout infancy, even over days, and these changes may influence neurovascular coupling, introducing noise, and potentially diminishing the reliability of functional connectivity estimates in infants (Kozberg and Hillman, 2016). Second, resting fMRI studies of infants are conducted during sleep (without sedation), whereas studies in adults and children are more commonly done while participants are awake. Different sleep states may be associated with distinct connectivity patterns, yet sleep state is seldom assessed or controlled for in infant fMRI research. Third, the increased water content of the infant brain, relative to a more mature brain, alters tissue contrast (e.g., T2-weighted images in infants provide clearer grey-white matter demarcation) and may make image processing more challenging, eroding measurement reliability. Fourth, functional networks in infants may be in a premature state, and potentially less stable, rendering their measurement less reliable.

Our primary objective in this paper is to investigate the short-term (within one MRI scanning session) test-retest reliability of functional connectivity measures in the newborn infant brain across four different studies, each conducted at a different site (***Table 1***). First, we investigate edge-level and subject-level test-retest reliability measured by ICCs from ROI-based whole-brain connectivity, as well as from two resting fMRI networks commonly investigated by ROI analysis – the default mode network (DMN) and the sensorimotor network (SMN). Second, we examine the influence on ICCs of run duration, study site, and scanner manufacturer (e.g., Philips and General Electric). Lastly, we test the spatial reproducibility from networks derived from independent component analysis (ICA), a data-driven method to assess functional connectivity that is distinct from ROI-based approaches. For these ICA-based analyses, we compare the ICA-derived network across runs (and term this, *spatial reproducibility*) as well as with canonical ICA-derived networks from adults (and termed this, *spatial similarity*).

**Table 1.** Demographics and MRI parameters by study

|  |  | Missing<sup>a</sup> | Study 1 | Study 2 | Study 3 | Study 4 | p-value |
| --- | --- | --- | --- | --- | --- | --- | --- |
| **Number of scans** |  |  | 43 | 23 | 27 | 26 |  |
| **PMA, mean (SD)** |  | 9 | 46.70<br>(4.63) | 42.30<br>(1.20) | 42.04<br>(0.90) | 49.53<br>(4.512) | <0.001 |
| **Sex, n (%)** | F | 8 | 19 (51.4) | 12 (54.5) | 14 (53.8) | 15 (57.7) | 0.991 |
|  | M |  | 18 (48.6) | 10 (45.5) | 12 (46.2) | 11 (42.3) |  |
| **Scanner** |  |  | GE 3T | GE 3T | Phillips 3T | GE 3T |  |
| **TR (sec)** |  |  | 2.0 | 2.2 | 2.3 | 2.0 |  |
| **Runs** |  |  | 2 | 5 | 1 | 1 |  |
| **Numb. frames/ run** |  |  | 216 | 102 | 210 | 210 |  |
| **Number of usable frames/run after scrubbing mean [min, max]** |  |  | 200<br>[185, 216] | 90<br>[76,102] | 179<br>[126,210] | 190<br>[153,210] |  |

PMA, postmenstrual age; M, male, F, female Study 1: Environmental influences on child health outcomes – Boricua Youth Study (ECHO-BYS) at the New York State Psychiatric Institute (NYSPI); Study 2: Serotonin system and neurodevelopment; Study 3, Brazil Babies Study; Study 4, ECHO-BYS at the University of Puerto Rico; <sup>a</sup>Missing PMA and/or sex occurred in Study 1 and 3.

## 2. METHOD

### 2.1 Datasets

We conducted this secondary data analysis from a sample of 239 infants acquired as part of four neurodevelopmental studies. After quality checking (see ****2.2**** for details), we included 119 infants in our analyses. ****Table 1**** summarizes the demographics for these 119 infants, including sex and postmenstrual age (PMA) at scan, as well as the scanning parameters used for each study. In the ****Supplementary materials,**** we provide a brief description of each study including their enrollment strategy.

### 2.2 Data analytic plan

As outlined in **Figure 1**, our data analytic plan consisted of four components: data curation, data processing, connectivity estimates, and test-retest reliability.

**Figure 1.**
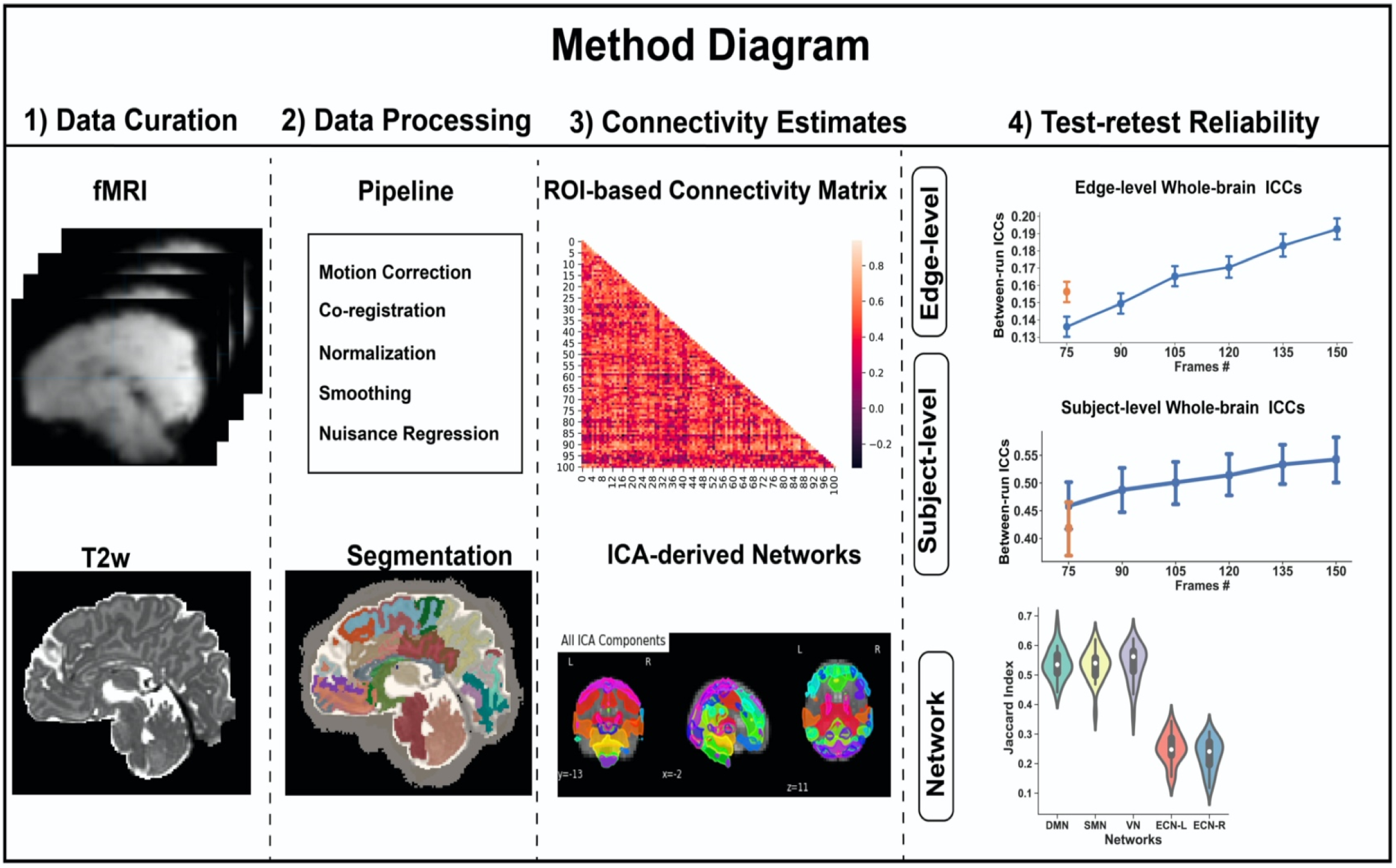

#### 2.2.1 Data Curation

We aggregated infant MRI datasets across the four neurodevelopmental studies yielding 239 infant scans. We conducted an initial quality assessment of the T2-weighted and resting fMRI scans by trained technicians who visually inspected the scans and removed those with incomplete data (e.g., no T2w scan and/or severe imaging artifacts, such as ghosting). A total of 90 out of 239 scans (~37.6%) were excluded on this basis. ***Supplementary Table 1*** indicates the number of scans excluded for each study.

#### 2.2.2 Data processing

##### 2.2.2.a. Scrubbing

For the resting fMRI scans, frames (or volumes) corrupted by high motion were marked as outliers and removed – a process called “scrubbing” (Mongerson et al., 2017). We adopted a common criterion for identifying outlier frames – a combination of frame-wise displacement (FD) and blood-oxygen level dependent (BOLD) data variance (DVARS). DVARS is an estimate of framewise changes in BOLD signal intensity and is calculated as the root-mean-squared variance of the temporal derivative of all brain voxel time series (Power et al., 2012). As suggested elsewhere (Power et al., 2013), frames were marked as outliers and removed if either of the following were true: FD◻>◻0.2◻mm or DVARS >3%.

We excluded entire functional runs if 1) there was only one run per subject, and the total scanning length after scrubbing was less than 5 mins; OR 2) there was more than one run per subject, but the run length after scrubbing was less than 2.5 mins. These criteria translated approximately to a combination of 2.5 minutes minimum per run after scrubbing and no more than 30-40% of scrubbed frames per run. Thirty scans were removed based on these criteria, leaving 119 usable scans for our analyses (see ***Supplementary Table 1***). The percentage of outlier frames within these 119 usable scans is summarized in ****Supplementary Figure 1****. We also compared head motion after scrubbing, based on frame-wise displacement (FD), across our four study sites and did not detect significant site differences (see ***Supplementary Figure 2***).

##### 2.2.2.b. Pre-processing functional data

After visual inspection and scrubbing, the functional scans were motion-corrected using MCFLIRT (Jenkinson et al., 2002), which transforms functional frames via rigid alignment and outputs an estimate of the six rigid motion parameters. We then implemented brain extractions (Smith, 2002), used a high-pass filter with a 100-second period cutoff value, and a low-pass spatial filter using FSL’s SUSAN, and a kernel of 4.5 millimeters for spatial smoothing (Smith and Brady, 1997). Finally, we scaled the mean signal intensity to 10,000.

##### 2.2.2.c. Nuisance regression

To reduce physiological noise due, for example, to cardiac and respiratory rhythms, and to minimize the influence of head motion, we regressed the whole-brain time course against the mean global signal, signal extracted from white matter (WM) and ventricular masks, as well as motion parameters (see Sect 2.2.2.b) and their derivatives. This approach to signal correction has been shown to provide a good trade-off between maintaining degrees of freedom and removing noise contamination (Ciric et al., 2017).

##### 2.2.2.d. T2-weighted segmentation and normalization

First, we used the Developmental Human Connectome Project automated pipeline for segmentation of T2w scans (Makropoulos et al., 2018). Second, for finer anatomical segmentation by lobe, we then further subdivided grey matter from the frontal, parietal, and occipital lobes with a custom algorithm that utilizes a warped segmentation from the individual subject to an AAL infant atlas (Shi et al., 2011). The estimated warp, or “normalization,” was carried out with Advanced Normalization Tools (ANTs). All normalization results were visually inspected to confirm accuracy. This modified segmentation pipeline generated 101 ROIs with whole-brain coverage, as listed in ***Supplementary Table 2***. Our finer segmentation pipeline code written in MATLAB is publicly available at GitHub: https://github.com/wangyuncolumbia/Infant_finer_segmentation.

##### 2.2.2.e. Co-registration of functional to structural and normalization to standard space

We calculated a mean functional volume and used this as the moving volume in an affine registration with the segmented structural scan, based on a normalized mutual-information (NMI) cost function in FLIRT (Smith et al., 2004). We then used this initial registration to guide a subsequent, and more accurate, boundary-based registration (BBR), also implemented with FLIRT, which utilizes the grey–white boundary to calculate the cost function (Greve and Fischl, 2009). Because averaging functional data can blur the grey–white boundary, we used an exemplar volume from the middle of the functional run for this BBR step. Lastly, we normalized the functional scans by applying the individual’s structural-to-template warp. To reduce the number of data interpolations to a single instance, we applied all linear transformations and the non-linear warp simultaneously through antsApplyTransforms (ANTs v2.1.0) using Lanczos interpolation.

### 2.3 Connectivity Estimates

#### 2.3.1 Region-of-interest (ROI)-based connectivity estimates

We derived a whole-brain ROI-to-ROI connectivity matrix based on our individualized segmentation of 101 ROIs – that is, we calculated a 101 x 101 connectivity matrix based on the Pearson correlation of the averaged time series fMRI data from each ROIs (***Supplementary Table 2***). We also generated ROI-to-ROI connectivity matrices for two resting fMRI networks – the DMN and the SMN, each with 10 ROIs (see ***Supplementary Table 2*** colored labels). We chose these two networks because they are often used in ROI-based functional connectivity analyses and prior research suggests that they begin forming early in neurodevelopment and thus may be more readily detected with resting fMRI techniques (Gao et al., 2015).

#### 2.3.2 Independent component analysis (ICA)-based connectivity

We used MELODIC to perform group-level independent component analysis (ICA) separately for each run and for each study (Smith et al., 2004). The number of components was set at 20 for our main analysis but varying the number of components from 15~75 yielded similar results (see ***Supplementary Figure 4***). To accurately identify and label resting fMRI networks extracted from the group ICA, we compared the components to well defined resting fMRI networks obtained using similar ICA procedures in adults (Smith et al., 2009). Using ITK-SNAP (Yushkevich et al., 2006) and Advanced Normalization Tools (ANTS), we registered and warped the 20 components derived from our infant group-ICA to a template infant atlas (Shi et al., 2011), and then similarly registered 20 components from an adult group-ICA (Smith et al., 2009) to the same template space. We then calculated the Jaccard index (see ***2.4*** for explanation of Jaccard index) between the infant and adult components and labeled the infant components based on their highest Jaccard index with a corresponding adult component (e.g., the infant ICA component showing the highest Jaccard index with the adult DMN was labeled the infant DMN component.) In addition, to examine spatial reproducibility across runs, we performed dual regression (Nickerson et al., 2017) to obtain subject-specific sets of spatial maps for each run and each subject. To binarize the data, we then thresholded the spatial maps at |z|>2.3. Lastly, to compare the spatial similarity of the networks identified across our study sites, we aggregated the functional data across runs from each study site and conducted a separate group ICA for each of the four study sites, then compared them using the Jaccard index.

### 2.4 Test-retest reliability

#### 2.4.1 Test-retest reliability from ROI-based analysis

To estimate the test-retest reliability for ROI-based connectivity, we calculated the Intraclass correlation coefficient (ICC [2,1]) (Chen et al., 2018; Noble et al., 2019). For Studies 1 and 2, which both included two or more runs, we compared the functional connectivity estimates between the two runs (Study 1) and between earlier runs to sequentially subsequent runs (Study 2), henceforth, referred to as “***between-run reliability***” (**Figure 2A**). Because Studies 3 and 4 included only one run, we could not calculate between-run reliability. Instead we compared functional connectivity estimates from the first half of the run with those from the second half – henceforth, referred to as “***within-run reliability***” (**Figure 2B**). For between-run and within-run reliability, we examined both edge-level and subject-level ICCs. To allow for site comparisons, we also calculated within-run reliability for Study 1. (We did not do this for Study 2 because the runs were only 75 frames, too short to estimate within-run reliability). The interpretation of the ICC is commonly categorized as follows: poor <0.4, fair 0.4–0.59, good 0.6–0.74, excellent ≥0.75 (Noble et al., 2019).

**Figure 2.**
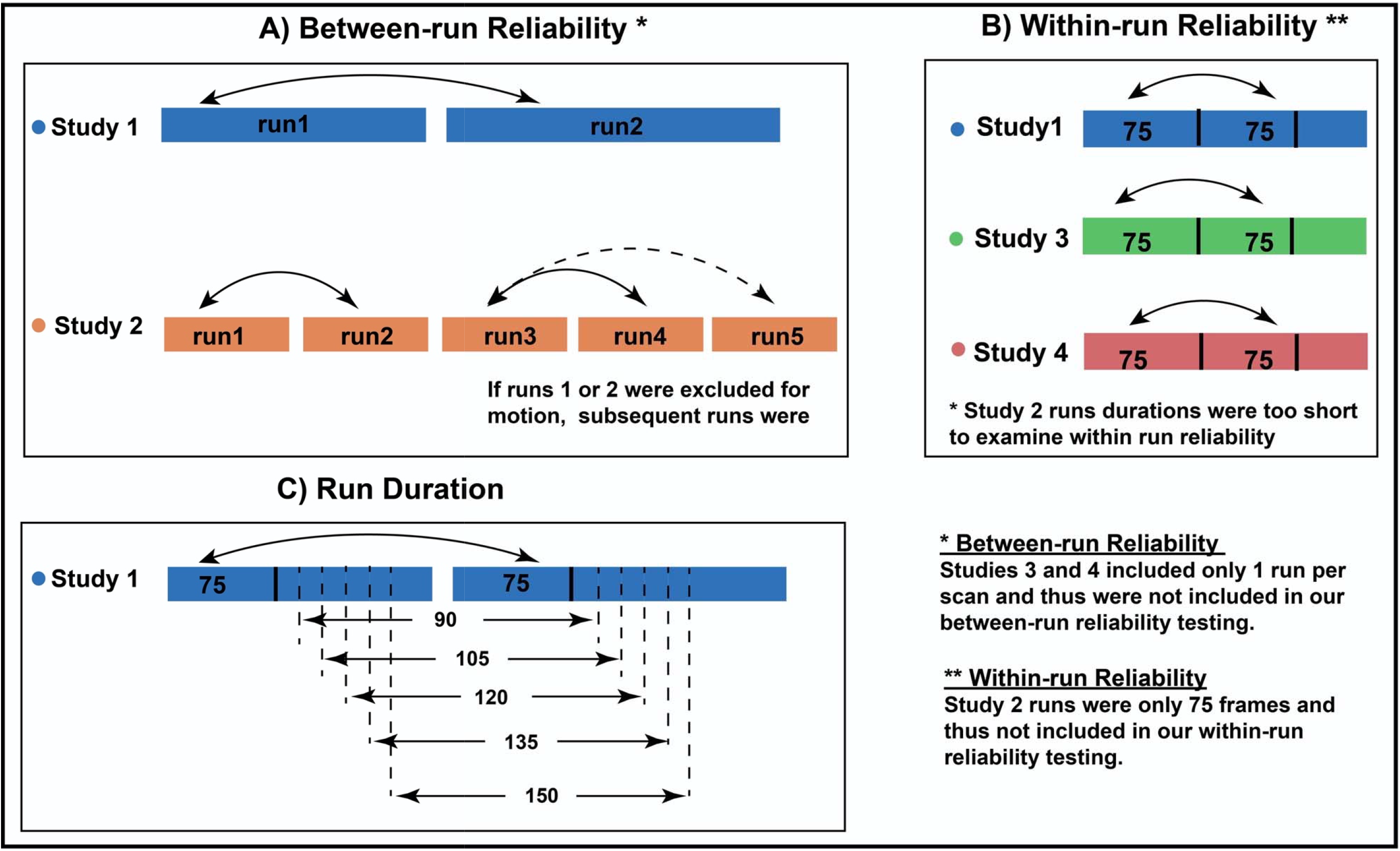

#### 2.4.2 Effects of run duration, site, and MRI manufacturer

We next aimed to examine the influence of run duration, study site, and MRI manufacturer on test-retest reliability. Because of collinearity between study site and MRI manufacturer, we examined the effect of study site and MRI manufacturer on “within-run reliability” in separate linear regression models, adjusting for age, sex, and head motion (as index by FD, see Sect 2.2). Furthermore, we conducted multiple linear regression to test for effects of run duration and study site (Studies 1 and 2 used the same type of MRI scanner) on “between-run reliability,” adjusting for age, sex, and head motion.

To construct different lengths of run duration, we used Study 1 because this study provided scans with two resting fMRI runs, each over 7 minutes in duration, which of our four studies provided the longest run durations. After scrubbing, we binned each scanning run into batches of frames, ranging from 75 frames to 150 frames in increments of 15 (**Figure 2C**). For example, for a scan with 150 frames or more in each of two runs, the data were binned into two sets of 75 frames, two sets of 90 frames, etc., up to two sets of 150 frames. The frames used were the initial frames for each run – for instance, in the previous example, the two sets of 75 frames were comprised of the first 75 frames from the first run, and the first 75 frames from the second run.

#### 2.4.3 Spatial reproducibility of ICA-derived networks

For our ICA-based analysis, we used the Jaccard index to calculate the spatial reproducibility of the derived networks across runs. The Jaccard index measures the similarity between two spatial network maps by computing the normalized amount of overlap; that is, the ratio of an intersection to the ratio of union of two spatial network maps. Jaccard Index ranges from 0 to 1; a high Jaccard index denotes high similarity of two spatial maps.

We limited our ICA-based analysis of spatial reproducibility to Study 1 because this study contained two runs allowing for comparison of the ICA-derived networks from the first run with those from the second. ICA estimates are not recommended for shorter runs (Glasser et al., 2018) and thus we excluded Study 2. Studies 3 and 4 were excluded because they contained only one run per infant.

#### 2.4.4 Spatial similarity of ICA-based networks with adult networks

We used the Jaccard index to evaluate the spatial similarity of ICA-derived networks from our infant studies with well-defined resting fMRI networks obtained using similar ICA procedures in adults (Smith et al., 2009). We conducted these comparisons for five well-defined resting fMRI networks: the default mode network (DMN), the sensorimotor network (SMN), the visual network (VN), and the left and right executive control network (ECN) and conducted these comparisons for each of our four study sites separately.

## 3. RESULTS

### 3.1 Test-retest reliability from ROI-based analysis

#### 3.1.1 Edge-level ICCs

We analyzed the reliability of all possible pairwise combinations of 101 ROIs across the whole brain, yielding 5,050 ICCs per scan. Between-run reliability was poor for runs of 75 frames (mean ICC = .146, 95% CI [.142, .151]) and poor for runs of 150 frames (mean ICC = .179, 95% CI [.175, .184]; see **Figure 3**). Within-run reliability was fair (mean ICCs = .41, 95% CI [.403, .410]; see ***Supplementary Figure 3***).

**Figure 3.**
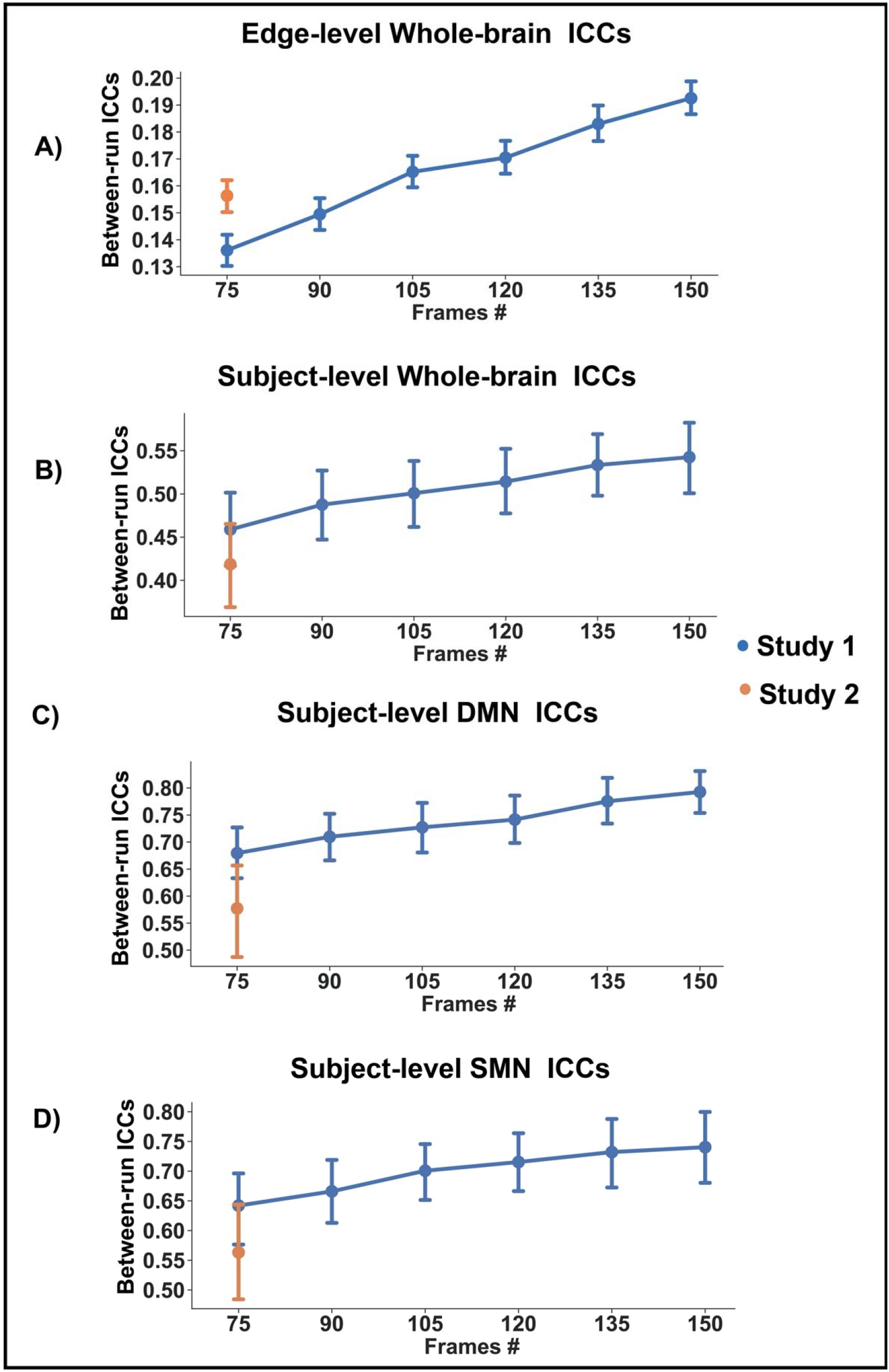

Regression analysis of between-run reliability showed a significant effect (F=81.95, p< 10^−10^) of run duration (t =12.64, p<10^−10^) and study site (t =−3.27, p=10^−3^). Linear regression on within-run reliability also confirmed a significant effect of study site (F =137.4, p< 10^−10^) and MRI scanner (F = 7.79, p< .005).

#### 3.1.2 Subject-level ICCs

##### 3.1.2.a Subject-level ICCs – Whole-brain analysis

Across the whole brain, between-run reliability was fair for runs of 75 frames (mean ICC = .43, 95% CI [.40, .47]) and fair-to-good for runs of 150 frames (mean ICC= .51, 95% CI [.46, .55]). Within-run reliability was good (mean ICCs=.63, 95% CI [.61, .65]). Regression analysis of between-run reliability showed a significant effect (F=10.32, p<10^−5^) of run duration (t=3.354, p=10^−3^) and study site (t =−3.17, p=.002). Neither significant effect of study site (F=0.25, p=0.62) nor MRI scanner (F=1.78, p=0.19) on within-run reliability was found.

##### 3.1.2.b Subject-level ICCs – Network-based analysis

For the DMN, between-run reliability was good for runs of 75 frames (mean ICC = .63, 95% CI [.59, .68]) and good-to-excellent for runs of 150 frames (mean ICC = .73, 95% CI [.69, .78]). Within-run reliability was also excellent (mean ICCs=.76, 95% CI [.73, .78]). Regression analysis of between-run reliability showed a significant effect (F=19.52, p< 10^−5^) of run duration (t=4.24, p< 10^−3^) and study site (t=−4.72, p< 10^−3^). Regression analysis of within-run reliability showed a significant effect of MRI scanner (F=14.50, p< 10^−3^) but not study site.

For the SMN, between-run reliability was fair-to-good for runs of 75 frames (mean ICC = .60, 95% CI [.55, .66]) and good for runs of 150 frames (mean ICCs ranges from .69, 95% CI [.63, .74]). Within-run reliability was good-to-excellent (mean ICCs = .72, 95% CI [.69, .74]). Regression analysis on between-run reliability showed a significant effect (F=11.93, p< 10^−5^) of run duration (t =3.22, p< 0.002), and study site (t=−3.77, p< 10^−3^). Regression analysis of within-run reliability showed a significant effect of MRI scanner (F=12.62, p< 10^−3^) but not study site.

### 3.2 Spatial reproducibility of ICA-derived networks

Using dual regression, we obtained subject-specific spatial maps for each of the two runs in Study 1 and identified five networks for comparison: DMN, SMN, VN, and the left and right executive control network (ECN). Based on the Jaccard index between runs, we found high spatial reproducibility of the DMN, SMN, and VN, but worse reliability for the left and right ECN (Jaccard indices: DMN=0.54, SMN=0.53, VN=0.54, left ECN=0.25, right ECN=0.23; **Figure 4B** presents the distribution of Jaccard indices stratified by network). We then compared the ICA-derived networks in infants with those from adults and found moderate spatial similarity between infants and adults for the DMN, SMN, and VN, but not for the left or right ECN (Jaccard indices ranged from 0.43 to 0.65 for DMN, SMN, and VN; but from 0.17 to 0.20 for left and right ECN; **Figure 4A**). Moreover, the Jaccard indices between the infant versus adult networks varied across sites; for example, for the DMN, it ranged from 0.64 for Study 1 to 0.39 for Study 2. As seen in **Figure 4C**, the precuneus and posterior cingulate – posterior hubs of the DMN – were consistently observed within the infant DMN across all four studies, however, the medial prefrontal cortex – an anterior hub of the DMN – was only encompassed within the DMN component in Studies 1 and 3. In ***Supplementary Figure 5***, we present comparisons of the SMN, VN, and the left and right executive control network (ECN) across our four study sites.

**Figure 4.**
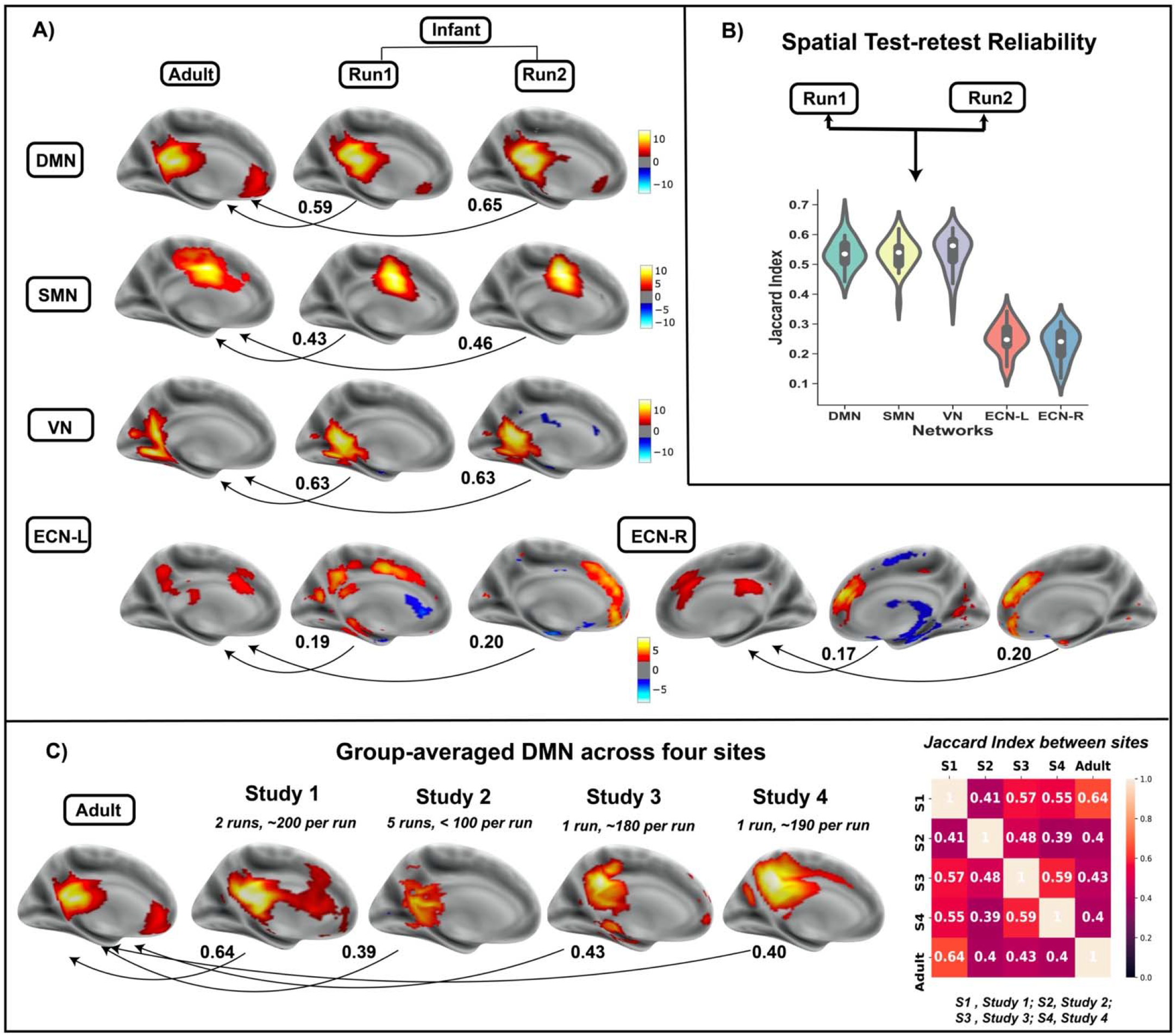

## 4. DISCUSSION

To our knowledge, this is the first study to investigate the short-term (intra-session) test-retest reliability of functional connectivity in infants across multiple sites and run durations. Three observations were noted. First, whereas individual connections (i.e. edge-level) exhibited poor reliability, subject-level reliability was moderate to high. Second, reliability improved as the number of frames increased, but reliability varied by study site and scanning manufacturer. Third, ICA-derived networks in infants exhibited moderate spatial reproducibility within scanning sessions and were spatially similar to adult networks; however, this was not true of all networks. Differences in spatial reproducibility across networks may indicate that networks differ in their maturational course with some developing earlier (e.g. DMN, SMN, and VN), and others later (e.g., left and right ECN), though this observation needs to be confirmed with longitudinal research.

Our reliability findings were largely in line with those reported in adult and child studies. For example, Pannunzi et al. examined the test-retest reliability of functional connectivity in 50 healthy adults and reported low reliability for single connections with an average edge-level ICC of 0.22 (Pannunzi et al., 2018). Even with long scan durations of 30 minutes (Birn et al., 2013), edge-level ICCs remained limited (~0.4). A recent meta-analysis of 25 resting fMRI studies encompassing over 2,000 participants concluded that individual connections have an average ICC of 0.29 (Noble et al., 2019). Conversely, when functional connectivity is examined at the subject level, reliability generally improves (Mejia et al., 2018; Mejia et al., 2015), with longer scanning lengths improving these results even further (Mejia et al., 2018). Recent advances, termed precision resting fMRI, suggest that high sampling rates coupled with extended scanning session (e.g. 60 + minutes), test-retest reliability can reach ICCs in the excellent range (0.8-0.9) (Gordon et al., 2017; Gratton et al., 2019). Additionally, the spatial reproducibility of ICA-derived networks have been consistently reported to be high in adults (Braun et al., 2012; Marchitelli et al., 2016). In sum, our reliability findings from infant resting fMRI data contrast poor edge-level reliability with good subject-level reliability and parallel the existing literature from adults.

Our reliability findings are simultaneously both reassuring and concerning. On the one hand, developmental neuroscientists should take some measure of comfort in knowing that subject-level measures from infant resting fMRI data can be estimated reliably with relatively short scanning durations (e.g. 5-10 minutes). On the other hand, our findings also suggest caution against reliance upon measures of individual functional connections. This is unfortunate because an individual functional connection is the most basic unit of analysis in resting fMRI research, and its interpretation compared to subject-level measures is more straightforward. Nonetheless, several subject-level measures are available and can be used to conduct statistical comparisons or to test brain-behavior correlations – for example, higher-order graph theory metrics (network density and clustering) (Braun et al., 2012; Cao et al., 2014; Termenon et al., 2016; Wang et al., 2019), and dynamic functional measures (Choe et al., 2017). Moreover, we found that reliability varied across resting fMRI networks derived from ICA, scanning sites and by scanning manufacturer. These last two points underscore the importance of data harmonization for multi-site studies, although site differences may persist nonetheless (Noble et al., 2017). Lastly, we did not find significant associations between reliability and infant age at scan, suggesting for example, that connectivity measures can be compared across infancy, albeit within the limited age range examined in our studies (postmenstrual age 39 – 59 weeks).

Limitations of our study are important to consider. First, our data quality procedures resulted in nearly 50% of scans being excluded. Less rigorous data quality procedures may impact reliability, and potentially reduce measurement validity. Moreover, this high level of data loss underscores the difficulty in obtaining high quality infant MRI data. Allowing ample time to prepare infants for the MRI scan (e.g. feeding and swaddling) and to fall asleep is imperative for successful data collection. Second, none of the four studies in our analysis used multiband imaging, which can significantly increase sampling rates. Increasing the rate of data acquisition, multiband imaging might improve the reliability of infant functional connectivity measures, although some research suggests that reliability is more affected by scan duration than sampling rate (Birn et al., 2013; Demetriou et al., 2018). Third, we used the Developmental Human Connectome Project automated pipeline (Makropoulos et al., 2018) to segment the infant brains in our study, and added additional parcellations to subdivide the frontal, parietal, and occipital lobes (see ***2.2.2***). Other approaches to infant brain segmentation are available (e.g. the newly released Infant FreeSurfer pipeline) and merit investigation. The approach to segmentation can influence the reliability of functional connectivity estimates – for example, adult resting MRI research indicates that more accurate segmentation improves functional connectivity measures (Salehi et al., 2020). To date, however, comparisons of tools for infant brain segmentation are scarce (Hashempour et al., 2019). Moreover, the infant brain is highly dynamic, increasing substantially in size and organization over development. As a result, for any given segmentation tool, its accuracy may exceed others at one time point in infancy, and yet become less accurate at subsequent time points. Brain segmentation with artificial intelligence (AI), by affording accuracy, speed, and flexibility, may offer a path forward, maintaining precision despite the dynamism of the infant brain. AI-based tools are still under development and will require validation (Wang, 2020).

In sum, our study may offer useful parameters for the use of infant resting fMRI as a methodology for neurodevelopmental research. For example, whereas caution should be taken against interpreting behavioral associations, or the predictive validity, of specific functional connections, developmental scientist may be on firmer ground when associating the functional organization of networks (e.g. DMN and SMN) with neurodevelopmental outcomes. Infant resting fMRI research is rapidly advancing, and judicious interpretation of its measures is essential to its ongoing success.

## Notes

### Competing Interest Statement

The authors have declared no competing interest.

